# Processing body dynamics drive non-genetic MEK inhibitors tolerance by fine-tuning KRAS and NRAS translation

**DOI:** 10.1101/2021.12.06.471470

**Authors:** Olivia Vidal-Cruchez, Victoria J Nicolini, Tifenn Rete, Roger Rezzonico, Caroline Lacoux, Julien Fassy, Karine Jacquet, Marie-Angela Domdom, Chloé Ventujol, Thierry Juhel, Barnabé Roméo, Jérémie Roux, Arnaud Hubstenberger, Bernard Mari, Baharia Mograbi, Paul Hofman, Patrick Brest

## Abstract

Overactivation of the Mitogen-activated protein kinase (MAPK) pathway is a critical driver of many human cancers. However, therapies targeting this pathway have proven effective in only a few cancers, as cancers inevitably develop resistance. Puzzling observations suggest that MAPK targeting fails in tumors due to early compensatory RAS overexpression, albeit by unexplained mechanisms. We identified a novel mechanism of drug tolerance to MEK inhibitors (MEKi) that involves Processing Bodies (PBs), a membraneless organelle (MLO). MEKi promoted translation of the oncogenes KRAS and NRAS, which in turn triggered BRAF phosphorylation. This overexpression, which occurred in the absence of neotranscription, depended on PB dissolution as the source of the *RAS* mRNA. Moreover, in response to MEKi removal, the process was dynamic as PBs rapidly reformed and reduced MAPK signaling. These results highlight a dynamic spatiotemporal negative feedback loop of MAPK signaling via *RAS* mRNA sequestration. Furthermore, we observed a phenotype with a low number of PBs along with strong KRAS and NRAS induction capacities. Overall, we describe a new intricate mechanism involving PBs in the translational regulation of essential cellular signaling pathways like MAPKs, paving the way for future therapies altering MLO and thereby improving targeted cancer therapies.

## Background

mRNAs are actively translated, repressed, stored, or degraded in response to environmental cues. These post-transcriptional pathways are critical for controlling cell fates that are altered under pathological conditions, such as cancer(1, 2). In this context, recent transcriptome-wide studies have shown that most RNAs have a restricted, dynamic and regulated subcellular localization. The key players in this process include two RNA clusters and their associated regulatory RNA-binding proteins (RBPs) known as constitutive processing bodies (PBs) and environmentally induced stress granules(3, 4). In contrast to organelles with a lipid bilayer membrane, membraneless structures are formed by a process known as liquid-liquid phase separation (LLPS), which confers a broad range of plasticity to these super-assemblies(5). Sequestration of mRNAs into these ribonucleoprotein granules is associated with translational repression, thus it uncouples mRNA expression from protein production and enables spatiotemporal control of gene expression(3). Moreover, PBs and stress granules act as reservoirs for silent mRNAs that can re-enter translation to adapt protein expression to the environment, and confer plasticity to the genetic program(3, 5–7). Although this post-transcriptional control is key to cellular adaptation to a volatile and stressful environment(1), few studies have investigated the role of PBs in the context of cancer progression and, in particular, as it relates to the emergence of drug-tolerant cells.

Mitogen-activated protein kinases (MAPKs) are ubiquitous signal transduction pathways that control cell fate by phosphorylating hundreds of substrates. The RAS-RAF-MEK-ERK pathway is altered in ∼40% of all human cancers, mainly through RAS oncogenic mutations (32%) and their downstream effector BRAF (∼10%). KRAS is mainly found mutated in pancreatic cancer (88%), lung cancer (30%) and colorectal adenocarcinomas (50%). NRAS and BRAF are altered in melanoma (17% and 55%, respectively), thyroid carcinoma (19% and 55%, respectively), and lung cancer (1% and 5%, respectively) (8, 9). Given the causative role of RAS-RAF-MEK-ERK pathway hyperactivation in tumorigenesis, several MEK, BRAF, and KRAS inhibitors (MEKi, BRAFi, and KRASi, respectively) have been developed in recent decades(10–12). Unfortunately, these treatments are inevitably associated with drug tolerance, acquired resistance and tumor recurrence(10, 13).

Although some resistance is directly attributable to the acquisition of somatic mutations, early drug resistance arose from plastic drug-tolerant or persister cells. Among these non-genetic early resistance events, upregulation of KRAS and NRAS proteins was suggested as a possible contributor in response to BRAFi or MEKi targeting downstream signalling pathway (14, 15), while the underlying mechanism remains unknown. Combining genetic and pharmacological approaches, we demonstrate that PBs are associated with drug tolerance through a DDX6-dependent negative feedback loop that fine-tunes the expression of KRAS and NRAS by controlling mRNA translation.

## Material and methods

### Cell culture

The A549, H1650, Mel501, BT549 were cultured according to the recommendations of the ATCC. A549 (human lung adenocarcinoma epithelial cell line, ATCC, number CCL-185) and Mel-501(melanoma epithelial cell line), were grown in Dulbecco’s Modified Eagle Medium (DMEM) supplemented with 10% fetal bovine serum (FBS). BT549 (Breast epithelial cell line, ATCC, number HTB-122) was grown in DMEM supplemented with 10% FBS and non-essential amino-acid (Thermofischer). H1650 (human lung adenocarcinoma epithelial cell line, ATCC, number CRL-5908) was grown in Roswell Park Memorial Institute (RPMI)-1640 medium supplemented with 10% FBS and 1mM sodium pyruvate. All the cells were maintained at 37°C in a 5% CO_2_ in a humidified incubator. All cells were maintained no more than one month and were identified using STR profile (Eurofins Genomics). For proliferation assay, 10 000 cells were seeded in six-well plates in triplicates and cells numbers were evaluated every day for 5 days using a Coulter counter (Villepinte, FRANCE).

### DDX6 GFP stable cell lines

The pPRIPu GFP-DDX6 plasmid used in this study were constructed as follows:

The pPRIPu CrUCCI vector(16) (kind gift of Dr F. Delaunay) was amplified with primer adaptors for AgeI and BamH1 restriction sites. pEGFP-C1_p54cp plasmids (kind gift of Drs D. Weil and M. Kress) were digested by AgeI and BamH1, and the respective resulting fragment was inserted in pPRIPU. The integrity of the entire sequence was confirmed by sequencing analysis. Briefly, replication-defective, self-inactivating retroviral constructs were used to establish stable A549 cell lines. Selection was performed with puromycin (10µg/ml). Cells were then sorted as a polyclonal population and used in subsequent experiments.

### siRNA transient transfection

Cells were plated at 200 000 cells/well in 6-well plates. After 24h, cells were transfected with negative Control siRNA (Silencer Pre-designed small interfering RNA) or with *DDX6*, *KRAS* or *NRAS* siRNA using JetPrime (PolyPlus) according to the manufacturer’s instructions as previously described(17). 48h after transfection, cells were lysed for RNA or protein analysis as described below.

### Immunoblotting

Immunoblotting was performed as described previously(17). Proteins were extracted from cells using Laemmli lysis buffer (12.5mM Na_2_HPO_4_, 15% glycerol, 3% sodium dodecyl sulfate [SDS]). The protein concentration was measured with the DC Protein Assay (BIO-RAD) and 30µg to 50µg of total proteins were loaded onto 7.5% or 12% SDS-polyacrylamide gels for electrophoresis and transferred onto polyvinylidene difluoride membranes (Millipore). After 1h of blocking with 5% bovine serum albumin or non-fat milk prepared in Phosphate-Buffered Saline (PBS)-0.1% Tween-20 buffer, the blots were incubated overnight at 4°C with antibodies (supp table). After 1h of incubation with a horseradish peroxidase-conjugated secondary antibody (1:3,000, Promega), protein bands were visualized using an enhanced chemiluminescence detection kit (Millipore) and and the Syngene Pxi4 imaging system (Ozyme). Western blot quantification was performed by Fiji freeware on unsaturated captured images.

### Isolation of RNA and quantitative real-time RT-PCR analysis

Total RNA extraction was performed using TRI-reagent (Sigma-Aldrich), followed by column based extraction as described previously(18). The RNA concentration was measured by NanoDrop 2000 (Thermo Fisher Scientific) and the integrity of RNA was analyzed by Bioanalyser (Agilent). For mRNA, the cDNA strand was synthesized from 500ng of total RNA. Quantification of *KRAS* and *NRAS* and *RPLP0* genes was measured by power-Sybr-green assays with the StepOne™ Real-time PCR System. The qPCR primers are referenced in the supplementary table.

### Immunofluorescence and confocal microscopy

Cells were grown to confluence and fixed in 4% paraformaldehyde for 20min. After fixation, cells were permeabilized with a solution containing 0.3% Triton X-100 for 5min. Then, cells were incubated with primary antibodies (supp table) overnight at 4°C in humidified chambers in a solution containing 0.03% Triton X-100, 0.2% gelatin and 1% BSA. Cells were washed and incubated with Alexa Fluor-conjugated secondary antibodies (1:500; Molecular Probes) for 1h at room temperature and mounted in ProLong Diamond Reagent with DAPI (Molecular Probes). Images were captured on a Zeiss LSM880 confocal microscope and analyzed with Fiji freeware.

### Polysome gradient

Subcellular fractionation. All steps of the subcellular fractionation were performed at 4°C. Cells (60-80×10 6 cells) with or without treatment were trypsinized and washed twice with 15 ml of ice-cold PBS by centrifugation. Cell pellets were resuspended in 1 ml of hypotonic Buffer composed of 40mM Tris-HCl, pH 7.5, 10mM KCl, 3mM MgCl_2_ and 0.2% Nonidet P-40 supplemented with 1mM DTT and 0,5mM phenylmethylsulfonyl fluoride (PMSF). After incubation for 15min on ice, cells were lysed in a Dounce homogenizer with B pestle. The efficiency of cell lysis in keeping the nuclei intact was verified by staining with trypan blue under an optical microscope. The cell lysates were spun at 900g for 5min to obtain the cytoplasm (supernatant) and nuclei (pellet) fractions.

Sucrose gradient fractionation. About 10 OD 260nm of cytosolic extracts were loaded to the top of a linear sucrose gradient (10–50%) made of Sucrose Gradient Buffer (25mM Tris-HCl, pH 7.5, 150mM NaCl, 12mM MgCl_2_, 1mM DTT) in Ultra-Clear ultracentrifugation tubes (Beckman Coulter). The cytosolic fraction samples were fractionated by ultracentrifugation for 2h 45min at 39,000rpm and at 4°C in a Beckman Optima ultracentrifuge with a SW41 Ti swinging bucket rotor at the following settings: acceleration 9 and deceleration 4. Following ultracentrifugation, the sucrose gradients were fractionated using a Foxy JR fraction collector and monitored using a UV light (254-nm wavelength) absorbance detector (Teledyne ISCO, UA-6), to obtain 12 to 14 fractions. After addition of 0,5mM CaCl_2_ and 0,2% SDS, the collected fractions were treated with proteinase K (50mg/ml, Sigma) for 30min at 40°C. Then, 10pg of an RNA spike-in was added to each fraction to serve as an internal calibrator of RNA extraction and detection. For this, we used an in vitro transcribed RNA that encodes the spike protein of SARS-CoV-2.

Total RNA was extracted by vigorous shaking with an equal volume of Tris pH 8,0 saturated phenol, chloroform, and isoamyl alcohol mixture (25:24:1, v/v/v) (Sigma-Aldrich), and phase separation was performed by centrifugation at 12,000g for 15min at 4°C. The upper aqueous phase was washed once with an equal volume of chloroform:isoamyl alcohol (24:1, v/v) (Sigma-Aldrich) by vigorous shaking and centrifugation at 12,000g for 15min at 4°C. The aqueous phase was transferred to a fresh tube and the RNA was precipitated overnight at −20°C by mixing with an equal volume of isopropanol, 2ml of glycoblue TM coprecipitant (Thermo Fisher Scientific) and 1/10 volume of 3M Na acetate pH 5.2. RNA was pelleted by centrifugation at 12,000g for 15min at 4°C.

### Statistical analysis

Quantitative data were described and presented graphically as medians and interquartiles or means and standard deviations. The distribution normality was tested with the Shapiro’s test and homoscedasticity with a Bartlett’s test. For two categories, statistical comparisons were performed using the Student’s t-test or the Mann–Whitney’s test. All statistical analyses were performed by the biostatistician using R.3.2.2 software and the Prism8.0.2 program from GraphPad software. Tests of significance were two-tailed and considered significant with an alpha level of p<0.05. (graphically: * for p<0.05, ** for p<0.01, *** for p<0.005).

## Results

### MEKi response promotes KRAS and NRAS translation

First, we confirmed the accumulation of NRAS and KRAS protein in response to MEKi, PD184352 (CI-1040), and trametinib in four different cell lines (A549, H1650, Mel501, BT549) regardless of the cancer type (lung, breast, or melanoma) and oncogenic drivers (KRASG12S, EGFR, BRAFV600E and PTEN, respectively) (Fig. 1A-C). To analyze whether both KRAS and NRAS proteins can activate downstream BRAF phosphorylation under these conditions (Fig. 1D-E), we used si*KRAS* and si*NRAS* either alone or in combination. Interestingly, we demonstrated that both KRAS or NRAS maintained BRAF phosphorylation in the presence of PD184352. Complete inhibition of BRAF phosphorylation was almost only achieved when both siRNAs were combined (Fig. 1D-E). Altogether, these results highlight that increasing protein expression of KRAS or NRAS was sufficient to induce phosphorylation of BRAF after MEK inhibition.

**Figure 1.**
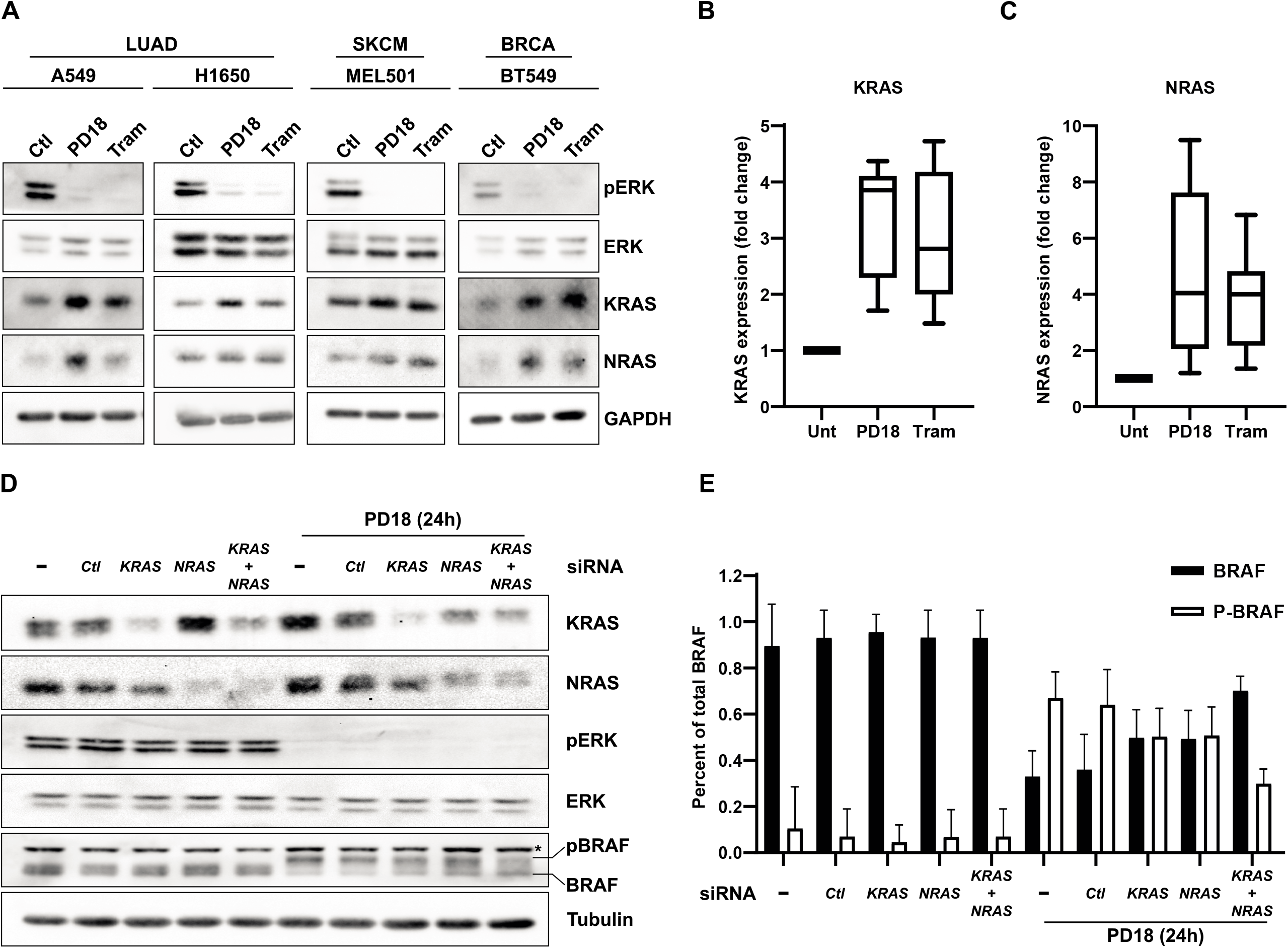
MEKi treatments induce potent KRAS and NRAS overexpression in cancer cells. **A.** Cancer cells were treated 24h with PD184352 (PD18) and trametinib (Tram) at 10µM and 10nM respectively. LUAD: Lung Adenocarcinoma; SKCM: Skin Cutaneous Melanoma; BRCA: Breast Cancer. Western blot analysis of the indicated proteins. pERK and pBRAF represent phosphorylated forms of ERK and BRAF respectively Results are representative of 3 independent experiments. **B-C.** Quantification of KRAS and NRAS expression by combining A549 (n=3), MEL501 (n=2), BT549 (n=2), and H1650 (n=1) biological replicates upon indicated treatments. **D.** A549 cells were transfected with the indicated siRNAs for 24h followed by 24h of treatment with PD184352 at 10µM (PD18). **E.** Quantification of BRAF forms in percent of total BRAF protein (n= 3 independent biological experiments).

In this context, we investigated the possible causes for this increase in KRAS and NRAS protein expression. At the transcriptional level, we showed that the mRNA levels of *KRAS* and *NRAS* were stable upon PD184352 treatment (Fig. 2A-B). Moreover, the PD184352 increase in KRAS and NRAS protein levels was maintained in the presence of the transcriptional inhibitor actinomycin D (Fig. 2C, suppl Fig. 1A). Altogether, these results argue for a posttranscriptional mechanism.

**Figure 2.**
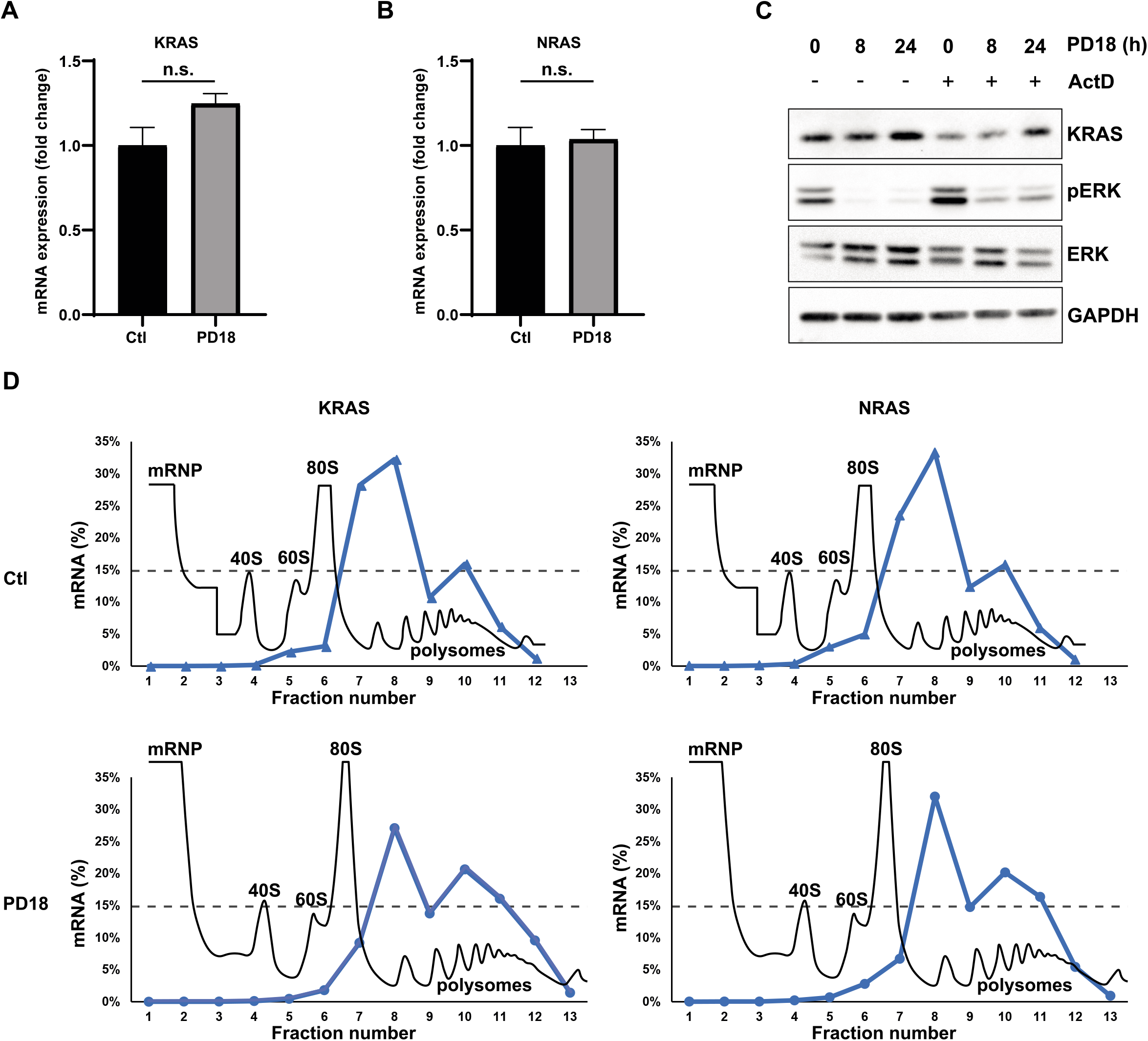
KRAS and NRAS overexpression is dependent on increased translation. **A-B.** A549 cells were treated with PD184352 (PD18) at 10µM and harvested after 24h. mRNA expression analysis of the indicated genes were analyzed by RT-qPCR using RPLP0 for normalization and untreated (Ctl) as reference (n=5 independent experiments). A Mann-Whitney test was performed for statistical analysis. (n.s.: non-significative) **C.** A549 cells were treated with PD184352 (PD18) at 10µM and Actinomycin D (ActD) at 1µg/mL and harvested at the indicated time. Western blot analysis of the indicated proteins (n=3 independent experiments). **D.** Polysome profiles (grey) of A549 cells treated 24h with PD184352 (PD18) at 10µM. One representative profile from two independent experiments is shown. RNA levels (Blue) of indicated transcripts in each polysomalfraction obtained by sucrose-gradient ultracentrifugation was quantified by qRT–PCR. Dotted-line represents baseline.

Using a ribosome profiling experiment, we showed that the mRNAs of *KRAS* and *NRAS* shifted from monosomes to heavy polysome fractions in the presence of PD184352 (Fig. 2D). To confirm an increase in translation, we treated A549 cells with the proteasome inhibitor MG132 to accumulate proteins by blocking their degradation. In cells treated with PD184352 and MG132, accumulation of KRAS was greater than with MG132 alone, confirming enhanced translation of *KRAS* mRNA in the presence of PD184352 (suppl. Fig. 1B-C). We conclude from these results that the rapid PD184352-induced NRAS and KRAS overexpression is due to an increase in translation rather than transcription.

### MEK inhibition promotes PB dissolution

Since it has been reported that *KRAS* and *NRAS* mRNAs are accumulated in PBs (3, 19), we hypothesized that protein overexpression of KRAS and NRAS is associated with the fact that the *KRAS* and *NRAS* pool of mRNA previously-stored in PBs has become available for translation. In this context, we analyzed the number and size of cellular PBs in different cancer cell lines. Strikingly, we observed a significant decrease in the size and number of PBs in response to PD184352 and trametinib in all cell types examined (Fig. 3A-F, Suppl. Fig. 2). This MEKi-induced decrease was most significant after 8h, showing a PB dissolution kinetics correlating with the time course of KRAS and NRAS overexpression. At this time point, MEKi had no effect on cell cycle progression, ruling out the possibility that PB dissolution was a secondary effect of cell cycle arrest (Fig. 3G-I, Suppl. Fig. 2). This overall decrease in the number of PB was not associated with lower DDX6 or LSM14A levels, two proteins required for PB formation (Fig. 3I), nor with the induction of stress granules (Suppl. Fig. 3).

**Figure 3.**
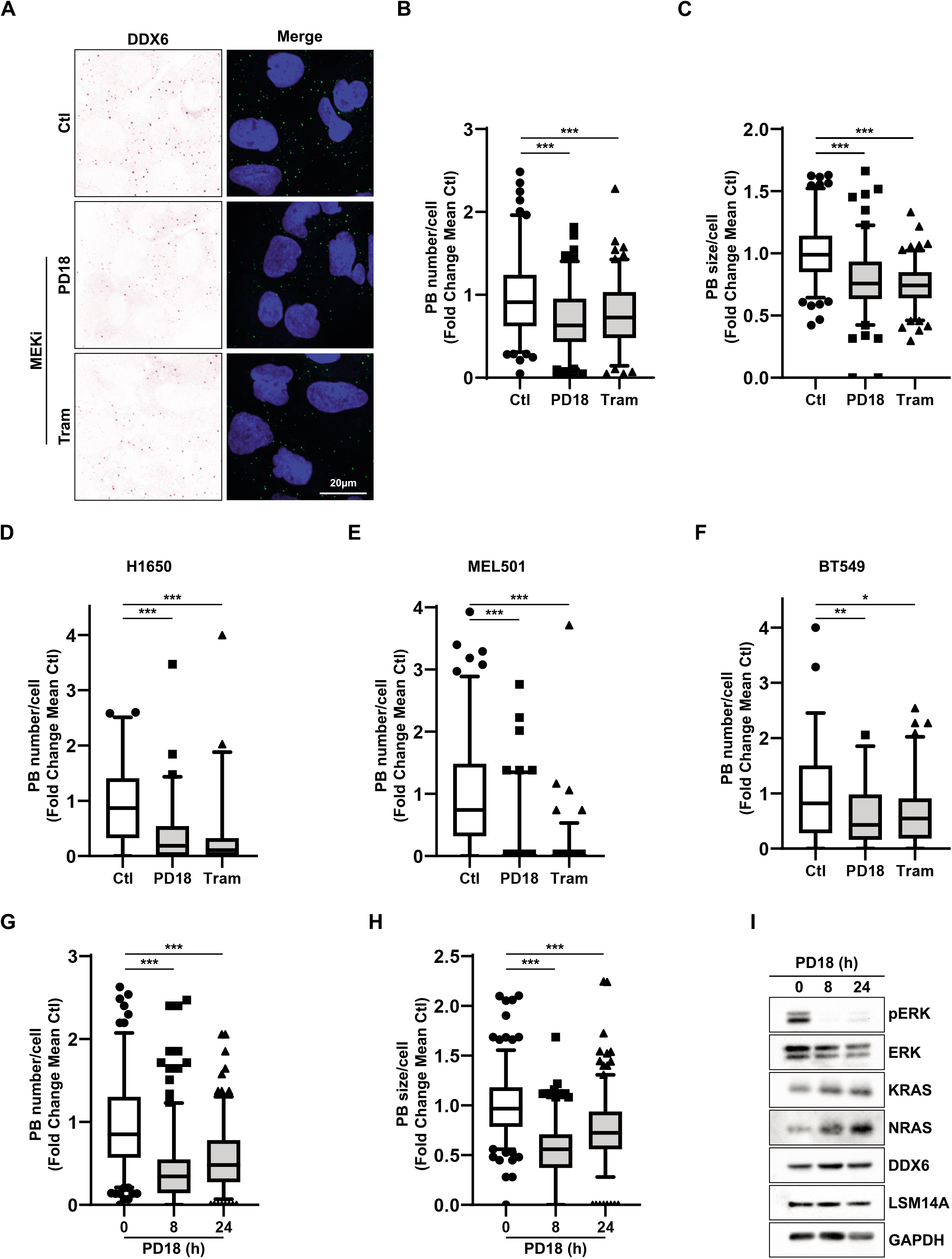
MEKi treatment is associated with a decrease in PB size and number. **A.** A549 cells were treated 24h with PD184352 (PD18) and trametinib (Tram) at 10µM and 10nM respectively Confocal analysis of PBs using anti-DDX6 antibodies (Inverted/green) with DAPI nuclear staining (blue). Results are representative of 3 independent experiments. **B-C** Quantification of indicated PB parameters of previous experiments by FIJI. Results are representative of 3 independent and merged experiments with a minimal quantification of 120 cells in total per conditions. A Mann-Whitney test was performed for statistical analysis. The p-value was indicated as *<0.05, **<0.01, ***<0.005. **D-F.** H1650, MEL501, BT549 cells were treated 24 h with PD184352 (PD18) and trametinib (Tram) at 10µM and 10nM respectively. Quantification of indicated PB parameters by FIJI. Results are representative of 3 independent merged experiments with a minimal quantification of 70 cells in total per conditions. A Mann-Whitney test was performed for statistical analysis. The p-value was indicated as *<0.05, **<0.01, ***<0.005. **G.H.I** A549 cells were treated with PD184352 (PD18) at 10µM and harvested at the indicated time. **G.H.** Quantification of indicated PB parameters by FIJI. Results are representative of 3 independent merged experiments with a minimal quantification of 220 cells in total per conditions. A Mann-Whitney test was performed for statistical analysis. The p-value was indicated as *<0.05, **<0.01, ***<0.005. **I.** Western blot analysis at the indicated time. pERK represents a phosphorylated form of ERK. Results are representative of 2 independent experiments.

Next, we analyzed the dynamics of the formation of PBs and the expression of KRAS. To this end, after treatment with PD184352 for 24h to dissolve PBs and induce overexpression of KRAS and NRAS, MEKi were removed, and cells were harvested at different time points (Fig. 4A). After PD184352 washout, strong activation of ERK was observed that persisted for 4h, along with stable expression of KRAS and NRAS and phosphorylation of BRAF (Fig. 4B, suppl Fig. 4). From 8 to 24h, a gradual decrease in the expression of KRAS and NRAS along with a decrease in BRAF phosphorylation was inversely correlated with an increasing number of PBs over time (Fig. 4B-D, suppl Fig. 4), indicating translational arrest. After removal of the drug, a growth rate comparable to that of untreated cells was re-established, demonstrating the rapid adaptability of cancer cells (Fig. 4E). Thus, our results provide evidence of two things. Firstly, they show that the high plasticity orchestrated by PBs helps to adjust the translational rate RAS through negative feedback loops. Secondly, they demonstrate that these KRAS and NRAS overexpression can trigger BRAF phosphorylation in the presence of MEKi. This PBs/RAS balance differentiate drug tolerance when compared with growth condition of cancer cells.

**Figure 4.**
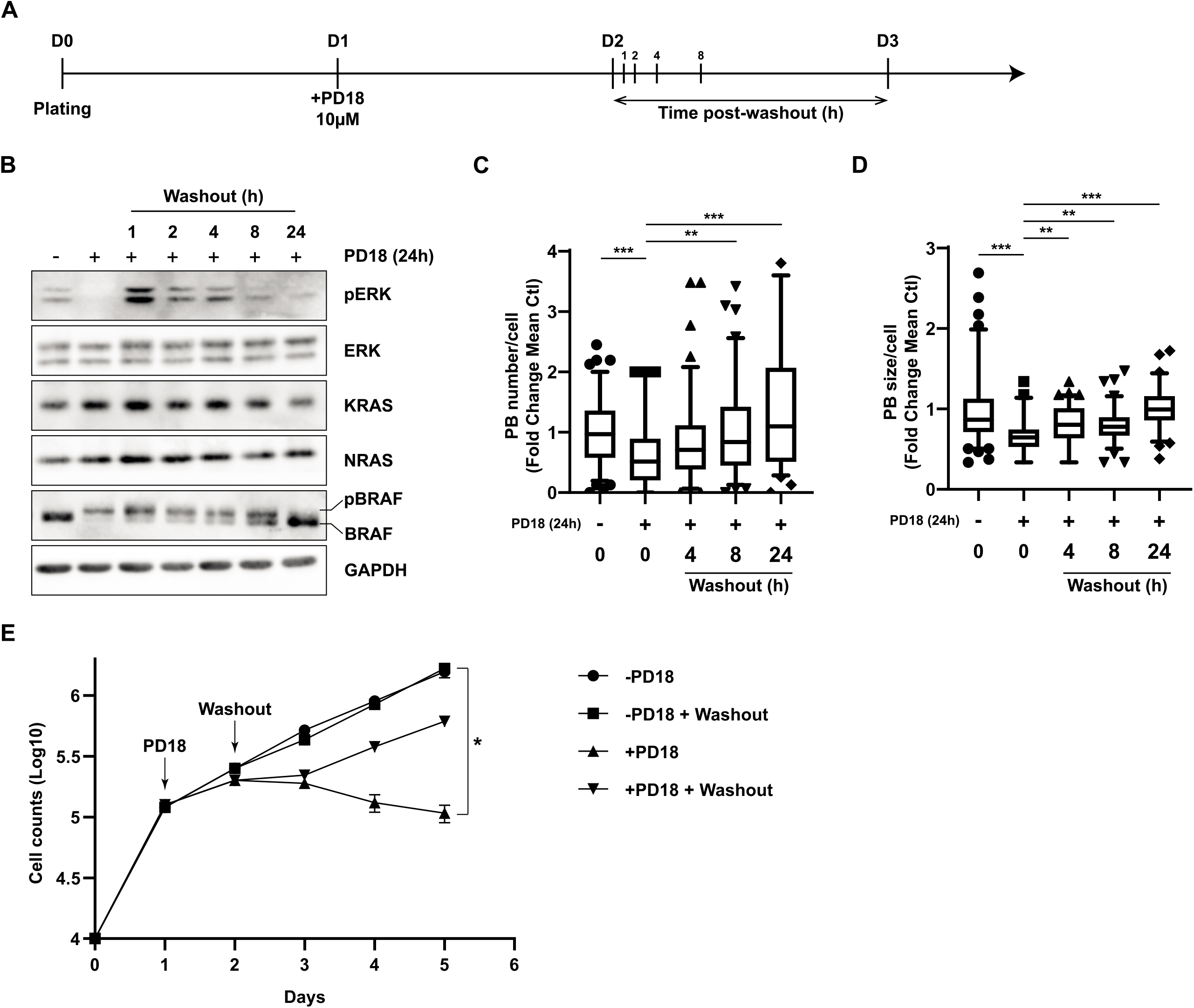
Dynamic regulation of PB formation and MAPK signaling. A549 cell were treated with PD184352 (PD18) at 10µM. After 24h MEKi was removed, and cells were harvested at the indicated time. **A.** Scheme of the experiments presented in panel B-D. **B.** Western blot analysis at the indicated time. pERK and pBRAF represents phosphorylated form of ERK and BRAF respectively. Results are representative of 2 independent experiments. **C.D.** Quantification by FIJI of PBs number and size respectively. Results are representative of 2 independent merged experiments with a minimal quantification of 60 cells in total per conditions. A Mann-Whitney test was performed for statistical analysis. For all experiments, the P-value was indicated as **<0.01; ***<0.005. **E**. Cell counts at indicated time. Results are representative of 2 experiments of 3 independent biological replicates.

Next, we tested whether PB dissolution and the associated of KRAS and NRAS overexpression could be corrected prolonged drug treatment. In drug-tolerant A549 cells, after tumor cells were treated with PD184352 at a dose of 2.5µM for 8 weeks, the drug was washed out and the drug-tolerant cells were then cultured for an additional 24h in the presence or absence of PD184352 at indicated concentrations. Figure 5 shows that activation of ERK after MEKi washout was identical to that observed in normal cells for 24h. Strikingly, the number and size of PBs were significantly lower than in the control condition, even in cells treated with a single dose of PD184352 (Fig. 5, Suppl. Fig. 5). Interestingly, at 10µM PD184352, the expression of KRAS was strongly induced in the resistant cells, along with a significant decrease in PBs (Fig.5, Suppl Fig.5). This overexpression was strong enough to maintain residual phosphorylation of ERK and overcame MEKi inhibition. As a sign of non-genetic resistance, these drug-tolerant cells restored a complete growth phenotype 5 days after MEKi removal (Suppl Fig. 5). Collectively, these results suggest that MEKi-induced PB dissolution is an early event in drug-tolerant cells and that this mechanism is established over time as a strategy to maintain KRAS and NRAS oncogenes expression to survive.

**Figure 5.**
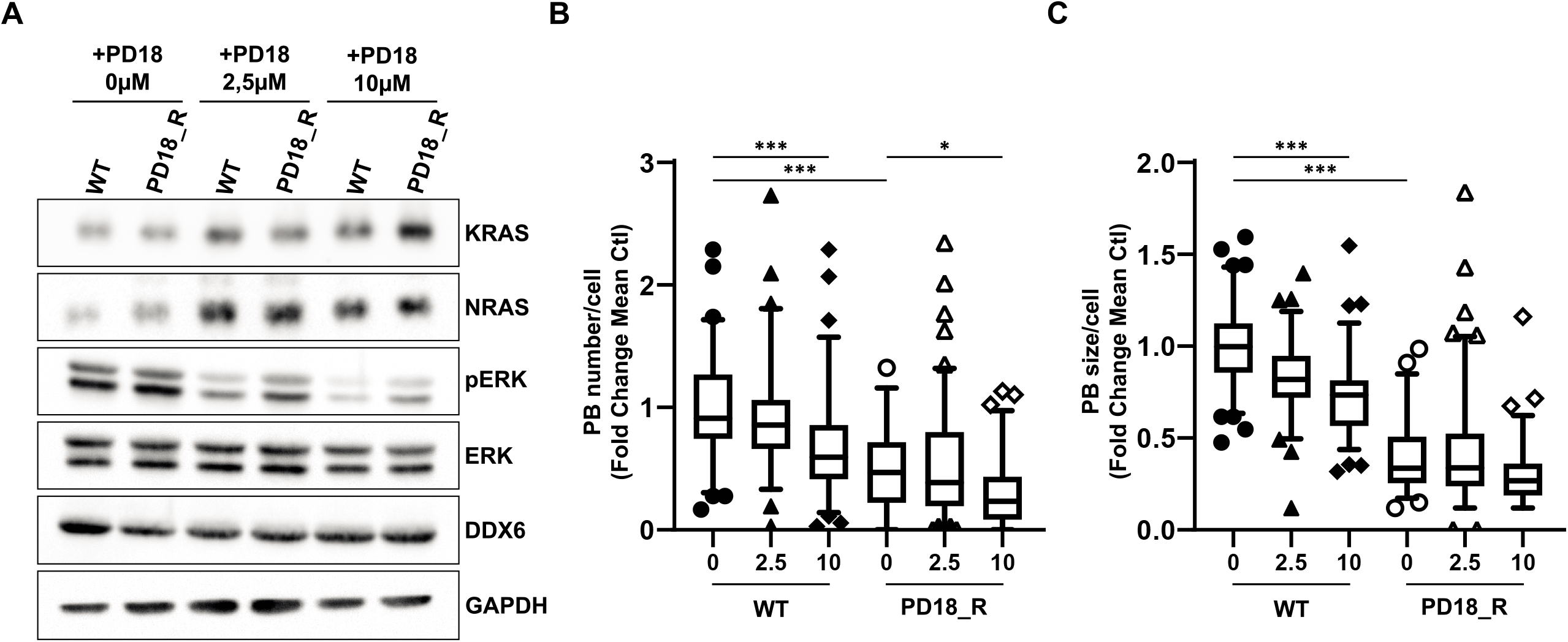
Resistant cells maintain a low-PB phenotype overtime. A549 wild type (WT) cells and 2.5µM PD18 resistant cells (PD18_R) were treated 24h with PD184352 (PD18) at 2.5µM or at 10µM. **A.** Western blot analysis at the indicated time. pERK represents phosphorylated form of ERK. Results are representative of 2 independent experiments. **B.C.** Quantification by FIJI of PBs number and size respectively. Results are representative of 2 independent merged experiments with a minimal quantification of 55 cells in total per conditions. A Mann-Whitney test was performed for statistical analysis. For all experiments, the P-value was indicated as *<0.05; **<0.01; ***<0.005.

### Essential PB components control KRAS and NRAS expression

To test for a causal role between the translational repression activities of PB components and KRAS and NRAS expression, we modulated the level of the RNA helicase DDX6 and essential PB component. Overexpression of GFP-tagged DDX6 increased the size and number of PBs. This significant increase in PBs correlated with decreased KRAS and NRAS expression (Fig. 6A-C, Suppl. Fig. 6). Notably, under these conditions, mRNA levels of *KRAS* and *NRAS* remained unchanged, indicating mRNA storage (Fig. 6D-E). Conversely, silencing of DDX6 resulted in complete dissolution of PBs, which was sufficient to trigger overexpression of KRAS and NRAS alone (Fig. 6F-G). Overall, these results demonstrated that PBs components play a critical role in mediating MEKi-induced RAS overexpression and the development of MEKi drug tolerance.

**Figure 6.**
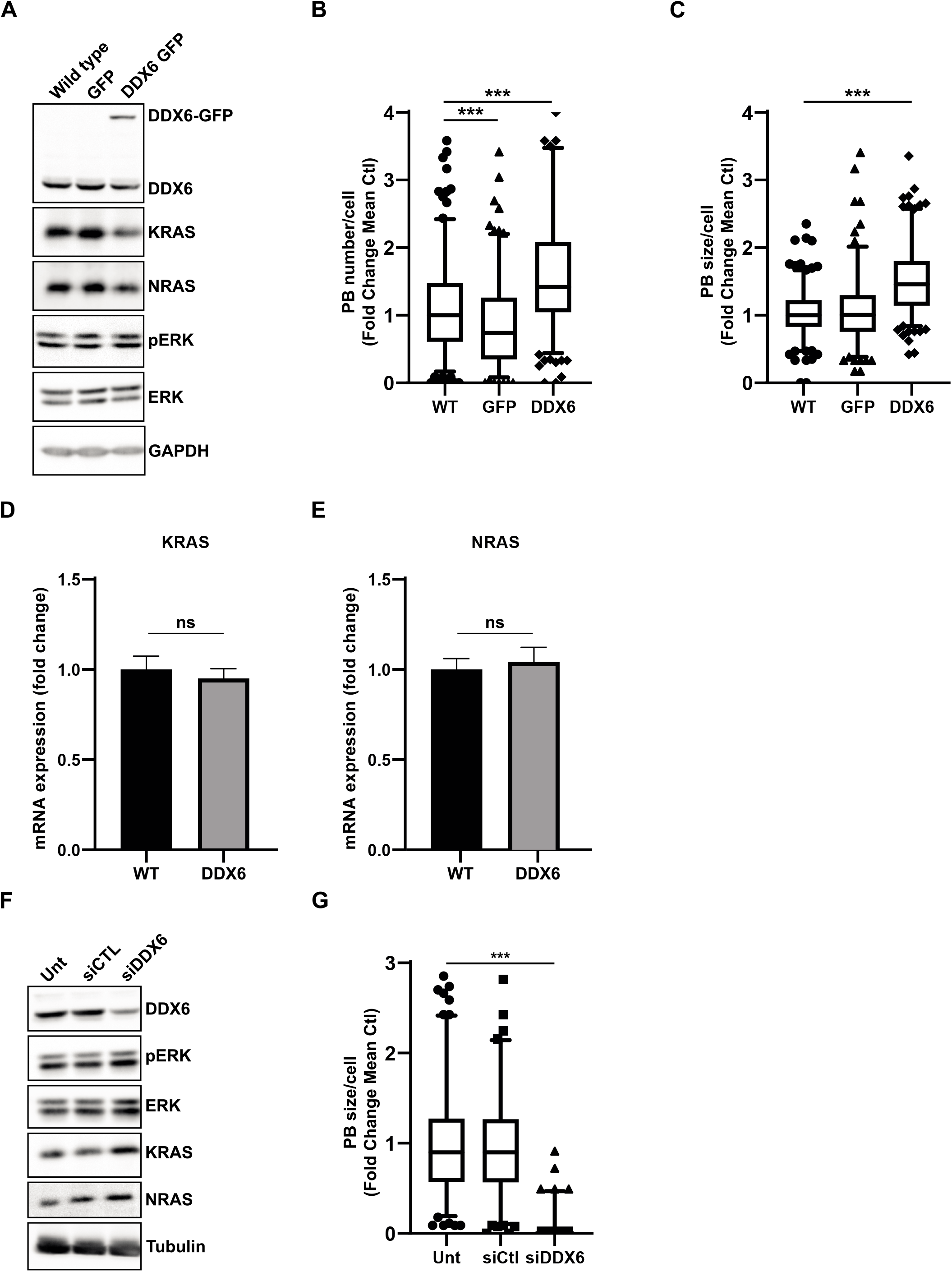
DDX6 expression controls KRAS and NRAS translation. **A-E.** Populations of A549 cells either wild type (WT), or overexpressing a green fluorescent protein (GFP), or a DDX6-GFP fused protein (DDX6). **A.** Western blot analysis at the indicated time. Results are representative of 2 independent experiments. **B.C.** Quantification by FIJI of PB number or size respectively. Results are representative of 3 independent merged experiments with a minimal quantification of 160 cells for each condition. A Mann-Whitney test was performed for statistical analysis**. D.E.** mRNA expression analysis of the indicated genes was analyzed by RT-qPCR using RPLP0 for normalization and untreated (WT) as reference. Results represent the combination of 3 independent experiments. A Mann-Whitney test was performed for statistical analysis. (n.s.: non-significative). For all experiments, the P-value was indicated as *<0.05; **<0.01; ***<0.005. **F.G**. A549 cells were transfected with a siControl (siCTL) or with a siDDX6 for 48h. **F.** Western blot analysis at the indicated time. Results are representative of 2 independent experiments. **G.** Quantification by FIJI of PB number. Results are representative of 3 independent merged experiments with a minimal quantification of 110 cells in total per condition. A Mann-Whitney test was performed for statistical analysis.

## Discussion

For many decades, studies have pointed to genetic mutations as a central mechanism for acquiring resistance to targeted therapies(20). However, there is a growing body of evidence that challenges this consensus and suggests that non-genetic heterogeneity and cell plasticity are actively involved in drug tolerance(21–23). Recently, new profiling techniques such as FATE -seq have been performed to unravel the non-genetic mechanisms of resistance(24). However, it is clear that genetic and non-genetic mechanisms of resistance or drug tolerance are often interrelated and not mutually exclusive(23, 25). Non-genetic resistance is due to the intrinsic plasticity of tumor cells, i.e., the ability to undergo transcriptional and epigenetic reprogramming in response to environmental challenges or to therapy. In this context, current therapeutic options for BRAFV600E/K patients include therapies targeting the MAPK pathway, which show remarkable efficacy in the first months of treatment(13). However, most patients treated with a combination of BRAF inhibitor (BRAFi) and MEK inhibitor (MEKi) inevitably relapse within a few months(13, 26). This relapse is associated with the presence of so-called persister, cancer stem, drug tolerant or resistant cells as reported in several studies, that harbor either a genetic or non-genetic program for survival(22, 27).

Here, we focused on the early events of drug tolerance. We demonstrated that within 8h, cells were able to establish overexpression of KRAS and NRAS that was independent of de novo transcription and inhibition of protein degradation. Interestingly, we observed that this overexpression persisted for weeks and eventually overcame MAPK inhibition. Drug-tolerant cells cultured in the presence of MEKi for a prolonged period exhibited a persistent decrease in PBs even 24h after treatment discontinuation, whereas restoration of PBs to normal levels occurred in drug-tolerant cells after a brief exposure to MEKi. Continuous versus intermittent BRAF and MEK inhibition in melanoma patients with BRAF V600E/K mutations was tested in a randomized, open-label phase 2 clinical trial (NCT02196181)(26). In this study, continuous administration resulted in a statistically significant improvement in progression-free survival after randomization compared to intermittent administration, suggesting that drug-tolerant cancer cells with increased plasticity may grow faster than long-term resistant cells after intermittent administration. Furthermore, by demonstrating that both KRAS and NRAS are equally important for maintaining BRAF activity, we have revealed novel ways to focus to replace MAPKi treatments, such as the development of resistance to KRAS G12C inhibitors in patients harboring either KRAS or NRAS mutations(11).

We have also shown that both PD184352 and trametinib can resolve MEKi. The role of the MAPK pathway is consistent with a previous study showing that MAPK3 (ERK1) was associated with PBs dissolution in the absence of a DDX6 decrease in the siRNA screening of 1,354 human genes(28). Our work highlights the fact that the activity rather than the level of ERK is critical for PBs dissolution. However, since ERKs can phosphorylate more than 200 intracellular targets, further studies are needed to determine whether the formation of PBs depends on the activity of ERK, either through direct targeting of PB components or indirectly through a partner of PBs.

Finally, we have shown that expression of the PB component DDX6, a helicase associated with miRNA-dependent repression of translation and PB storage, controls KRAS and NRAS translation. These results are consistent with previous studies showing accumulation of *KRAS* and *NRAS* mRNAs in PBs(3, 19), which likely depend on multiple *let-7* miRNA response elements on both 3’UTR(29). Moreover, several features of *KRAS* and *NRAS* mRNAs (i.e., the length of 5.3kb and 4.3kb, respectively, and the low GC content of their coding region or 3’UTR mRNA, 38% and 44%, respectively) may also contribute to their targeting of PBs(30). Previous studies have shown that KRAS is poorly translated compared to HRAS due to its enrichment in genomically underrepresented or rare codons(31, 32). In our study, we observed that either MEKi-induced dissolution or downregulation of DDX6 were associated with strong translation of *KRAS* and *NRAS* mRNAs and overexpression of the corresponding proteins. These results suggest that rare codons, often associated with the presence of an AU-rich third codon, favor the recruitment of mRNA repressors due to slow translational processing, which, however, can be overcome under certain conditions.

Overall, these results reveal a novel negative feedback loop involving PBs in the translational control of *KRAS* and *NRAS* mRNAs (Suppl Fig. 7). The type of regulation of the MAPK pathway that we have described here should pave the way for therapies that avert early resistance and achieve drug-tolerant cells before relapse.

## Ethics approval and consent to participate

Not applicable

## Consent for publication

Not applicable

## Availability of data and materials

Database sharing does not apply to this article as no RNA datasets were generated or analyzed during the current study. Additional files containing raw data experiments and links to https://idr.openmicroscopy.org/ for confocal microscopy collection of images will be provided for final revision and publication.

## Authors’ contributions

OVC and PB supervised and wrote the manuscript. OVC, VJN, TR, RR, CL JF, KJ, MAD, CV, TJ, and BR contributed substantially to define the concept and design the study; they acquired, analyzed, and interpreted of the data. JR, AH, BMa, BMo, and PH substantively revised the manuscript. AH, BMa, BMo, PH, and PB supported the work through fundings. All authors read and approved the final manuscript.

## Acknowledgments

Our thanks to the UCA Office of International Scientific Visibility for comments on the English version of the manuscript. Authors would like to thank Dr Julien Cherfils and Ludovic Cervera (Cytomed facility), Frederic Brau (IMPC) and Nadir Djerbi (PICMI) for technical help, Dr Dominique WEIL, Dr Michel Kress, and Dr Franck Delaunay for kindly providing plasmids.

## Abbreviations

MAPK: Mitogen-activated protein kinase
MEKi: MEK inhibitors
PBs: Processing Bodies
RBP: RNA binding proteins
LLPS: Liquid Liquid phase separation
DMEM: Dulbecco’s Modified Eagle Medium
FBS: fetal bovine serum
RPMI: Roswell Park Memorial Institute
siRNA: small interfering RNA
SDS: Sodium Dodecyl sulfate
PBS: Phosphate-Buffered Saline
BSA: Bovine serum albumin
PD18: PD184352
LUAD: Lung Adenocarcinoma
SKCM: Skin Cutaneous Melanoma
BRCA: Breast Cancer
ActD: Actinomycin
D Tram: Trametinib

---

**Supplementary Figure 1.**

**A.** A549 cells were treated with PD184352 (PD18) at 10µM and Actinomycin D (ActD) at 1µg/mL and harvested at the indicated time. Quantification of Western blot of KRAS (n=3 independent experiments). **B-C.** A549 cells were pretreated with MG132 at 2.5µM and treated with PD184352 (PD18) at 10µM and harvested at the indicated time. B. Western blot analysis at the indicated time. pERK represents phosphorylated form of ERK. **C.** Quantification of western blot.

**Supplementary Figure 2.**

**A.** H1650, **B.** MEL501 and **C.** BT549 cells were treated 24h with PD184352 (PD18) or trametinib (Tram). **D.E.** A549 cells were treated with PD184352 (PD18) at 10µM and harvested at indicated time. **A-D.** Confocal analysis of PBs using anti-DDX6 antibodies (inverted color/green) with DAPI nuclear staining (blue). **E.** Cell cycle distribution was analyzed by flow cytometry. Results represent the merge of 3 independent experiments.

**Supplementary Figure 3.**

A549 cells were treated 24h with PD184352 (PD18) at 10µM and 20min with Sodium Arsenite (NaAsO_2_) at 0.5mM. Confocal analysis of PBs using anti-DDX6 (Inverted/green) and stress granule anti-G3BP1 (Inverted/red) antibodies respectively with DAPI nuclear staining (blue).

**Supplementary Figure 4.**

A549 cells were treated with PD184352 (PD18) at 10µM, after 24h MEKi was removed, and cells were harvested at the indicated time. **A-B.** Western blot quantification of KRAS, NRAS, and pERK at the indicated time, Results represent the merge of 2 independent experiments. **C.** Confocal analysis of PBs using anti-DDX6 (Inverted/green) antibodies with DAPI nuclear staining (blue).

**Supplementary Figure 5.**

**A**. A549 wild type (WT) cells and 2.5µM PD18 resistant cells (PD18_R) were treated 24h with PD184352 (PD18) at indicated concentrations. Confocal analysis of PBs using anti-DDX6 (Inverted/green) and LSM14A (Inverted/Red) antibodies with DAPI nuclear staining (blue). **B.** Western blot quantification of KRAS and NRAS at the indicated time, Results represent the merge of 2 independent experiments **C**. Resistant cells (PD18_R) were cultured in the absence or presence of PD184352 (2.5µM) and counted at indicated time. Results are representative of 2 independent biological triplicates.

**Supplementary Figure 6.**

**A-B.** A549 wild type (WT) cells, overexpressing a green fluorescent protein (GFP) and a DDX6-GFP fused protein (DDX6). **A.** Western blot quantification of KRAS and NRAS at the indicated time, Results represent the merge of 2 independent experiments. **B.** Confocal analysis of PBs using anti-LSM14 (Inverted/green) antibodies with DAPI nuclear staining (blue). **C-E.** A549 cells were transfected with a siControl (siCTL) or with a siDDX6 for 48h. **C-D.** Western blot quantification of KRAS and NRAS at the indicated time. Results represent the merge of 2 independent experiments. **E.** Confocal analysis of PBs using anti-LSM14 (Inverted/green) antibodies with DAPI nuclear staining (blue).

**Supplementary Figure 7.**

Negative feedback loop under ERK1/2 phosphorylation is associated with formation of Processing bodies (PBs). PBs control *KRAS* and *NRAS* mRNA translation. PB dissolution under MEKi treatment is associated with enhanced *KRAS* and *NRAS* mRNA translation

## Supplementary Table

### List of reagents

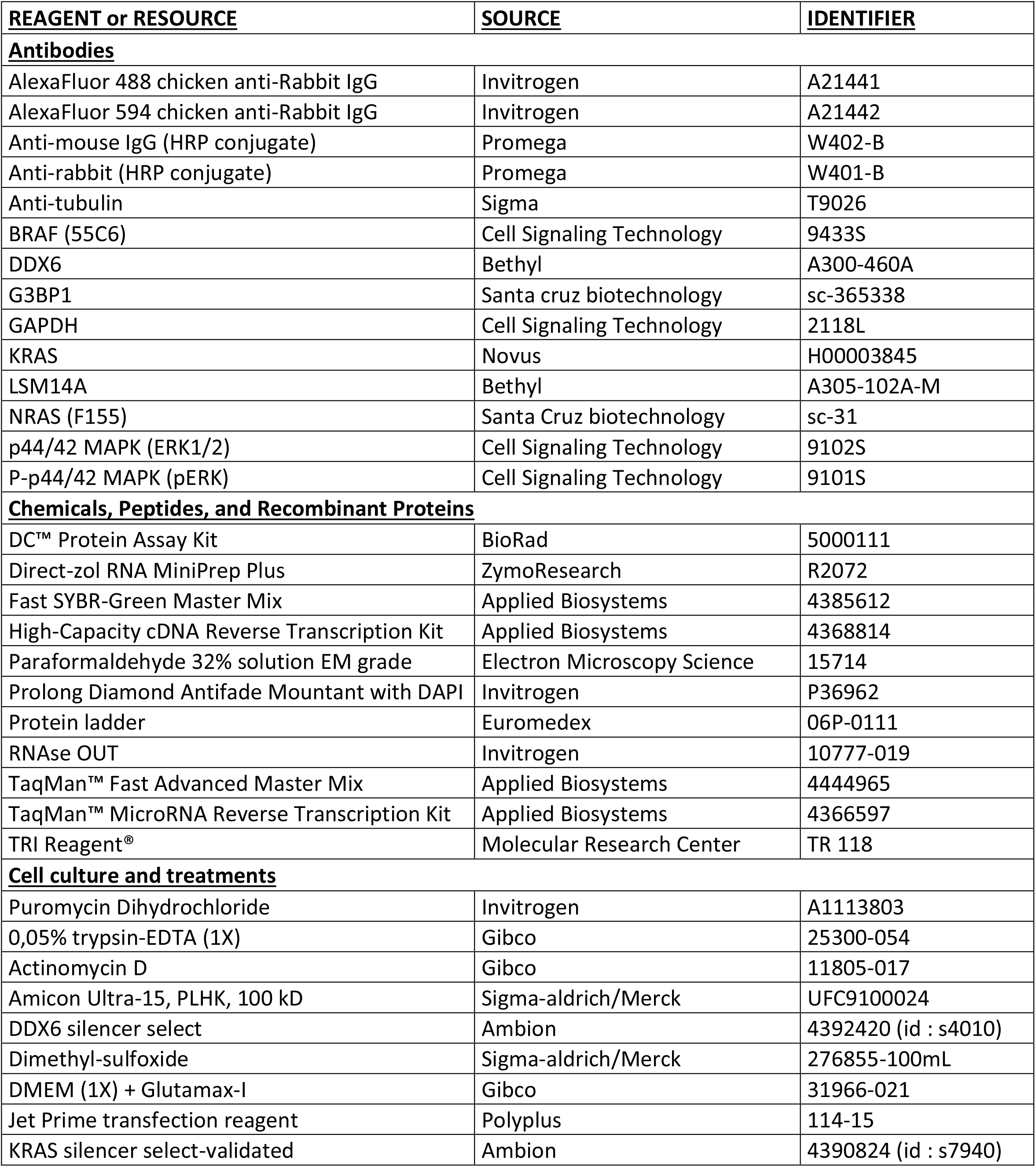
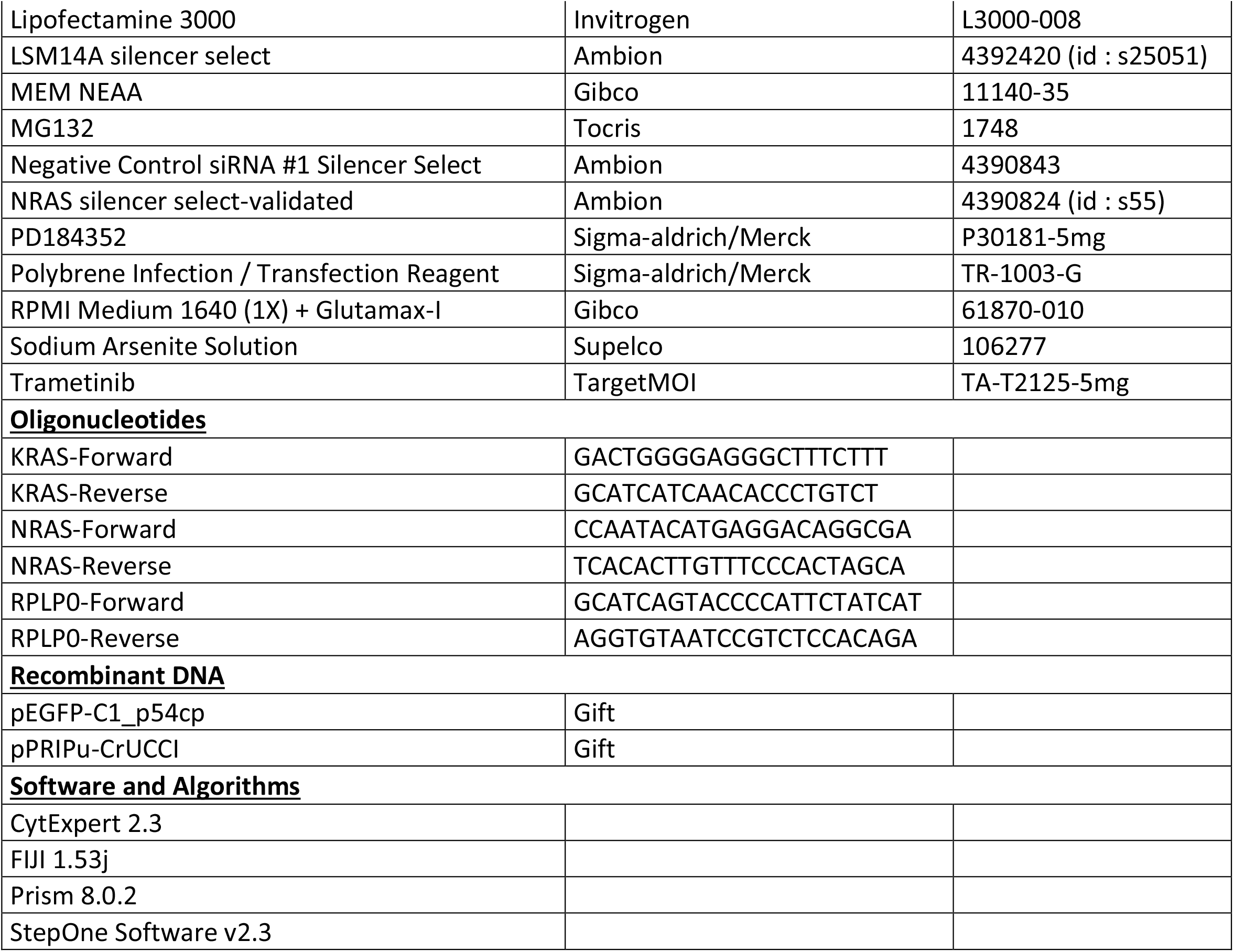

## References

1. Anderson, P., Kedersha, N. and Ivanov, P. (2015) Stress granules, P-bodies and cancer. Biochimica et Biophysica Acta (BBA) - Gene Regulatory Mechanisms, 1849, 861– 870.

2. Masuda, K. and Kuwano, Y. (2019) Diverse roles of RNA-binding proteins in cancer traits and their implications in gastrointestinal cancers. Wiley Interdisciplinary Reviews: RNA, 10, 1–18.

3. Hubstenberger, A., Courel, M., Bénard, M., Souquere, S., Ernoult-Lange, M., Chouaib, R., Yi, Z., Morlot, J.B., Munier, A., Fradet, M., et al. (2017) P-Body Purification Reveals the Condensation of Repressed mRNA Regulons. Molecular Cell, 68, 144–157.e5.

4. Khong, A., Matheny, T., Jain, S., Mitchell, S.F., Wheeler, J.R. and Parker, R. (2017) The Stress Granule Transcriptome Reveals Principles of mRNA Accumulation in Stress Granules. Molecular Cell, 68, 808–820.e5.

5. Banani, S.F., Lee, H.O., Hyman, A.A. and Rosen, M.K. (2017) Biomolecular condensates: Organizers of cellular biochemistry. Nature Reviews Molecular Cell Biology, 18, 285– 298.

6. Liu, J., Valencia-Sanchez, M.A., Hannon, G.J. and Parker, R. (2005) MicroRNA-dependent localization of targeted mRNAs to mammalian P-bodies. Nature cell biology, 7, 719–23.

7. Pitchiaya, S., Mourao, M.D.A.A., Jalihal, A.P., Xiao, L., Jiang, X., Chinnaiyan, A.M., Schnell, S. and Walter, N.G. (2019) Dynamic Recruitment of Single RNAs to Processing Bodies Depends on RNA Functionality. Molecular Cell, 74, 1–13.

8. Prior, I.A., Hood, F.E. and Hartley, J.L. (2020) The frequency of ras mutations in cancer. Cancer Research, 80, 2669–2974.

9. Anguera, G. and Majem, M. (2018) BRAF inhibitors in metastatic non-small cell lung cancer. Journal of Thoracic Disease, 10, 589–592.

10. Awad, M.M., Liu, S., Rybkin, I.I., Arbour, K.C., Dilly, J., Zhu, V.W., Johnson, M.L., Heist, R.S., Patil, T., Riely, G.J., et al. (2021) Acquired Resistance to KRAS G12C Inhibition in Cancer. New England Journal of Medicine, 384, 2382–2393.

11. Tanaka, N., Lin, J.J., Li, C., Ryan, M.B., Zhang, J., Kiedrowski, L.A., Michel, A.G., Syed, M.U., Fella, K.A., Sakhi, M., et al. (2021) Clinical Acquired Resistance to KRAS G12C Inhibition through a Novel KRAS Switch-II Pocket Mutation and Polyclonal Alterations Converging on RAS–MAPK Reactivation. Cancer Discovery, 10.1158/2159-8290.cd-21-0365.

12. Smith, L.K., Sheppard, K.E. and McArthur, G.A. (2021) Is resistance to targeted therapy in cancer inevitable? Cancer Cell, 39, 1047–1049.

13. Aldea, M., Andre, F., Marabelle, A., Dogan, S., Barlesi, F. and Soria, J.C. (2021) Overcoming resistance to tumor-targeted and immune-targeted therapies. Cancer Discovery, 11, 874–899.

14. Wang, Y., Van Becelaere, K., Jiang, P., Przybranowski, S., Omer, C. and Sebolt-Leopold, J. (2005) A role for K-ras in conferring resistance to the MEK inhibitor, CI-1040. Neoplasia, 7, 336–347.

15. Ambrosini, G., Khanin, R., Carvajal, R.D. and Schwartz, G.K. (2014) Overexpression of DDX43 mediates MEK inhibitor resistance through RAS upregulation in uveal melanoma cells. Molecular Cancer Therapeutics, 13, 2073–2080.

16. Feillet, C., Krusche, P., Tamanini, F., Janssens, R.C., Downey, M.J., Martin, P., Teboul, M., Saito, S., Lévi, F.A., Bretschneider, T., et al. (2014) Phase locking and multiple oscillating attractors for the coupled mammalian clock and cell cycle. Proceedings of the National Academy of Sciences of the United States of America, 111, 9828–9833.

17. Zangari, J., Ilie, M., Rouaud, F., Signetti, L., Ohanna, M., Didier, R., Roméo, B., Goldoni, D., Nottet, N., Staedel, C., et al. (2016) Rapid decay of engulfed extracellular miRNA by XRN1 exonuclease promotes transient epithelial-mesenchymal transition. Nucleic acids research, 45, 4131–4141.

18. Lassalle, S., Zangari, J., Popa, A., Ilie, M., Hofman, V., Long, E., Patey, M., Tissier, F., Belléannée, G., Trouette, H., et al. (2016) MicroRNA-375/SEC23A as biomarkers of the *in vitro* efficacy of vandetanib. Oncotarget, 7.

19. Pillai, R.S., Bhattacharyya, S.N., Artus, C.G., Zoller, T., Cougot, N., Basyuk, E., Bertrand, E. and Filipowicz, W. (2005) Inhibition of translational initiation by let-7 microRNA in human cells. Science, 309, 1573–1576.

20. Chen, Z., Fillmore, C.M., Hammerman, P.S., Kim, C.F. and Wong, K.-K. (2014) Non-small-cell lung cancers: a heterogeneous set of diseases. Nature Reviews Cancer, 14, 535–546.

21. De Conti, G., Dias, M.H. and Bernards, R. (2021) Fighting drug resistance through the targeting of drug-tolerant persister cells. Cancers, 13, 1–15.

22. Marin-Bejar, O., Rogiers, A., Dewaele, M., Femel, J., Karras, P., Pozniak, J., Bervoets, G., Van Raemdonck, N., Pedri, D., Swings, T., et al. (2021) Evolutionary predictability of genetic versus nongenetic resistance to anticancer drugs in melanoma. Cancer Cell, 39, 1135–1149.e8.

23. Marine, J.C., Dawson, S.J. and Dawson, M.A. (2020) Non-genetic mechanisms of therapeutic resistance in cancer. Nature Reviews Cancer, 20, 743–756.

24. Meyer, M., Paquet, A., Arguel, M.-J., Peyre, L., Gomes-Pereira, L.C., Lebrigand, K., Mograbi, B., Brest, P., Waldmann, R., Barbry, P., et al. (2020) Profiling the Non-genetic Origins of Cancer Drug Resistance with a Single-Cell Functional Genomics Approach Using Predictive Cell Dynamics. Cell Systems, 11, 367–374.e5.

25. Vendramin, R., Litchfield, K. and Swanton, C. (2021) Cancer evolution: Darwin and beyond. The EMBO Journal, 40, 1–20.

26. Algazi, A.P., Othus, M., Daud, A.I., Lo, R.S., Mehnert, J.M., Truong, T.G., Conry, R., Kendra, K., Doolittle, G.C., Clark, J.I., et al. (2020) Continuous versus intermittent BRAF and MEK inhibition in patients with BRAF-mutated melanoma: a randomized phase 2 trial. Nature Medicine, 26, 1564–1568.

27. Oren, Y., Tsabar, M., Cuoco, M.S., Amir-Zilberstein, L., Cabanos, H.F., Hütter, J.C., Hu, B., Thakore, P.I., Tabaka, M., Fulco, C.P., et al. (2021) Cycling cancer persister cells arise from lineages with distinct programs. Nature, 596, 576–582.

28. Berchtold, D., Battich, N. and Pelkmans, L. (2018) A Systems-Level Study Reveals Regulators of Membrane-less Organelles in Human Cells. Molecular Cell, 72, 1035–1049.e5.

29. Johnson, S.M., Grosshans, H., Shingara, J., Byrom, M., Jarvis, R., Cheng, A., Labourier, E., Reinert, K.L., Brown, D. and Slack, F.J. (2005) RAS is regulated by the let-7 microRNA family. Cell, 120, 635–647.

30. Courel, M., Clément, Y., Bossevain, C., Foretek, D., Vidal Cruchez, O., Yi, Z., Bénard, M., Benassy, M.-N., Kress, M., Vindry, C., et al. (2019) GC content shapes mRNA storage and decay in human cells. eLife, 8.

31. Fu, J., Dang, Y., Counter, C. and Liu, Y. (2018) Codon usage regulates human KRAS expression at both transcriptional and translational levels. Journal of Biological Chemistry, 293, 17929–17940.

32. Lampson, B.L., Pershing, N.L.K., Prinz, J.A., Lacsina, J.R., Marzluff, W.F., Nicchitta, C. V., MacAlpine, D.M. and Counter, C.M. (2013) Rare codons regulate KRas oncogenesis. Current Biology, 23, 70–75.

