## Supplementary figures and images for "Processing body dynamics drive non-genetic MEK inhibitors tolerance by fine-tuning KRAS and NRAS translation"

### Supplementary files

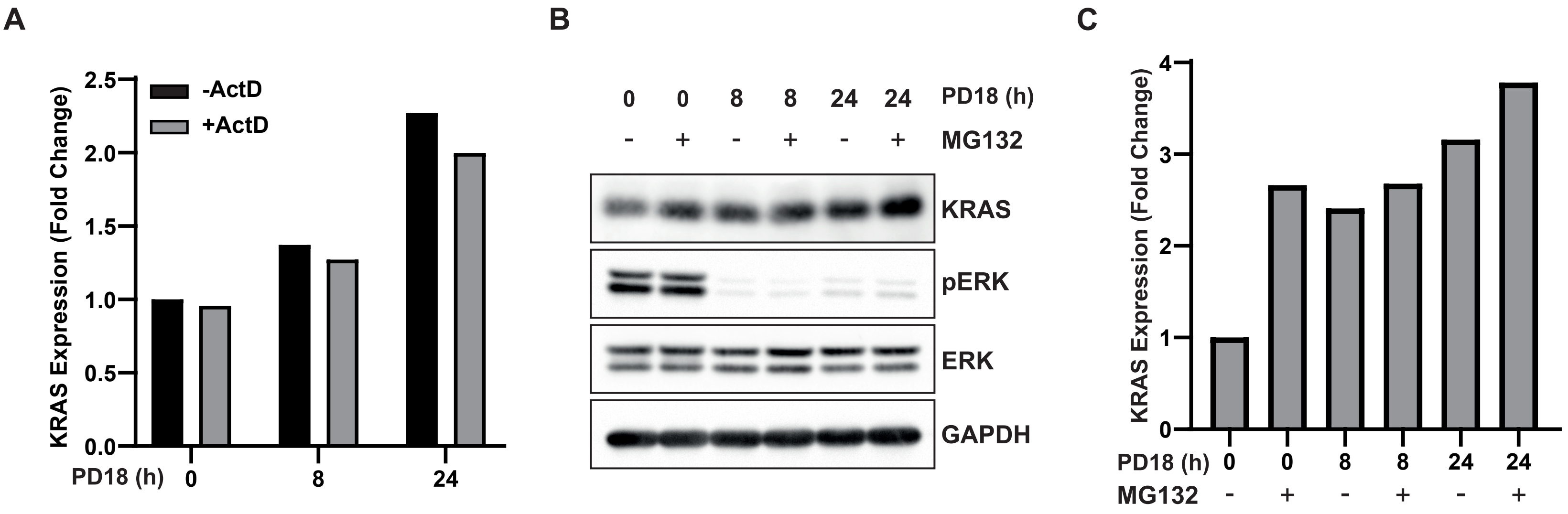


**Suppl. Figure 1**

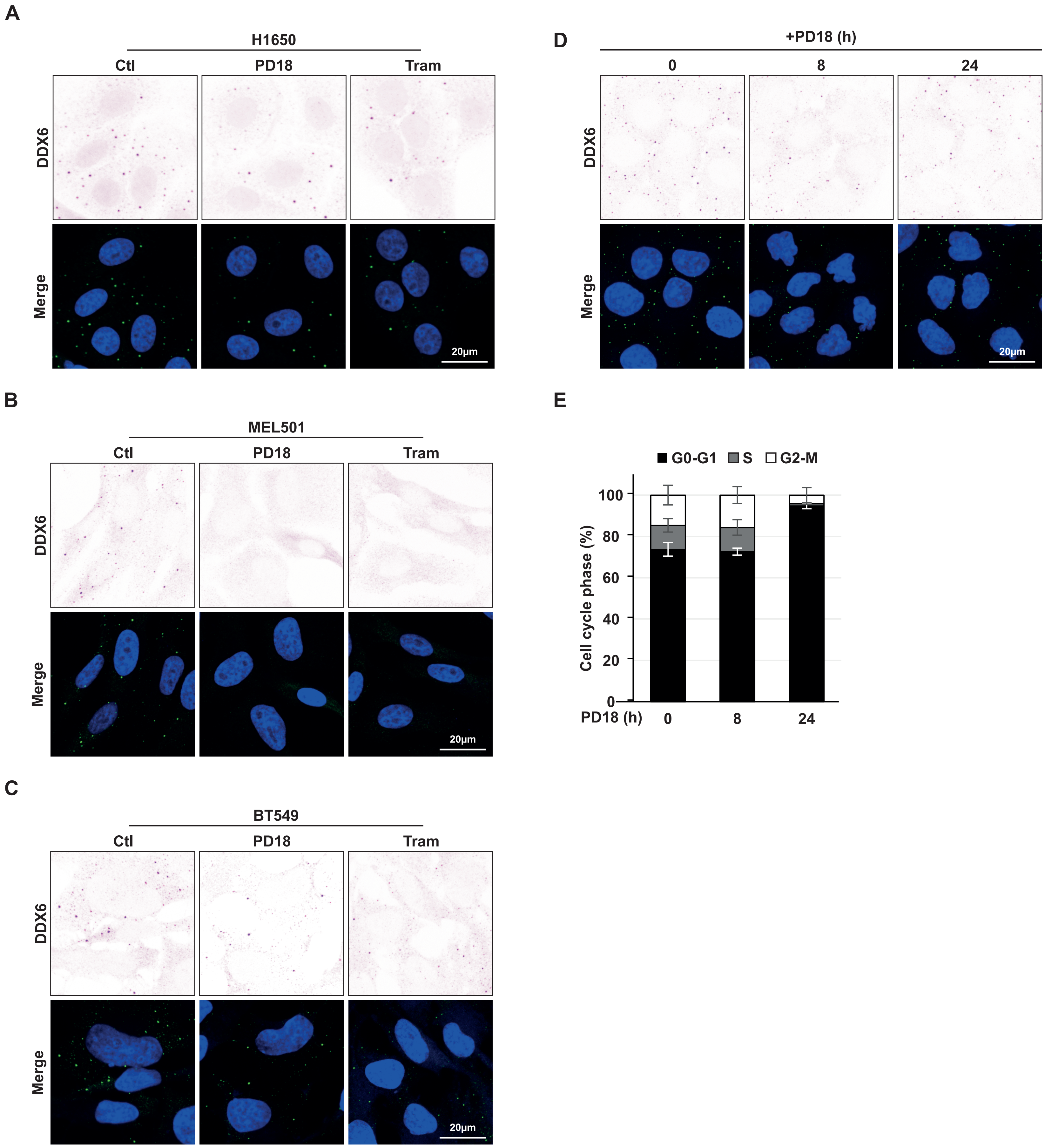


**Supp. Figure 2**

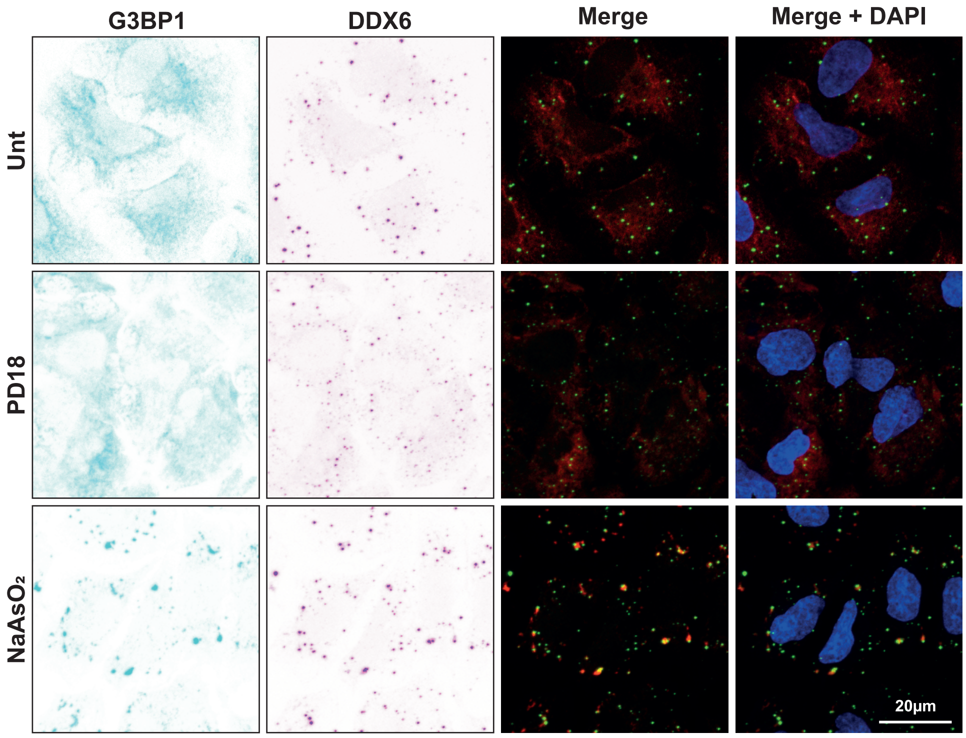


#### Supp. Figure 3

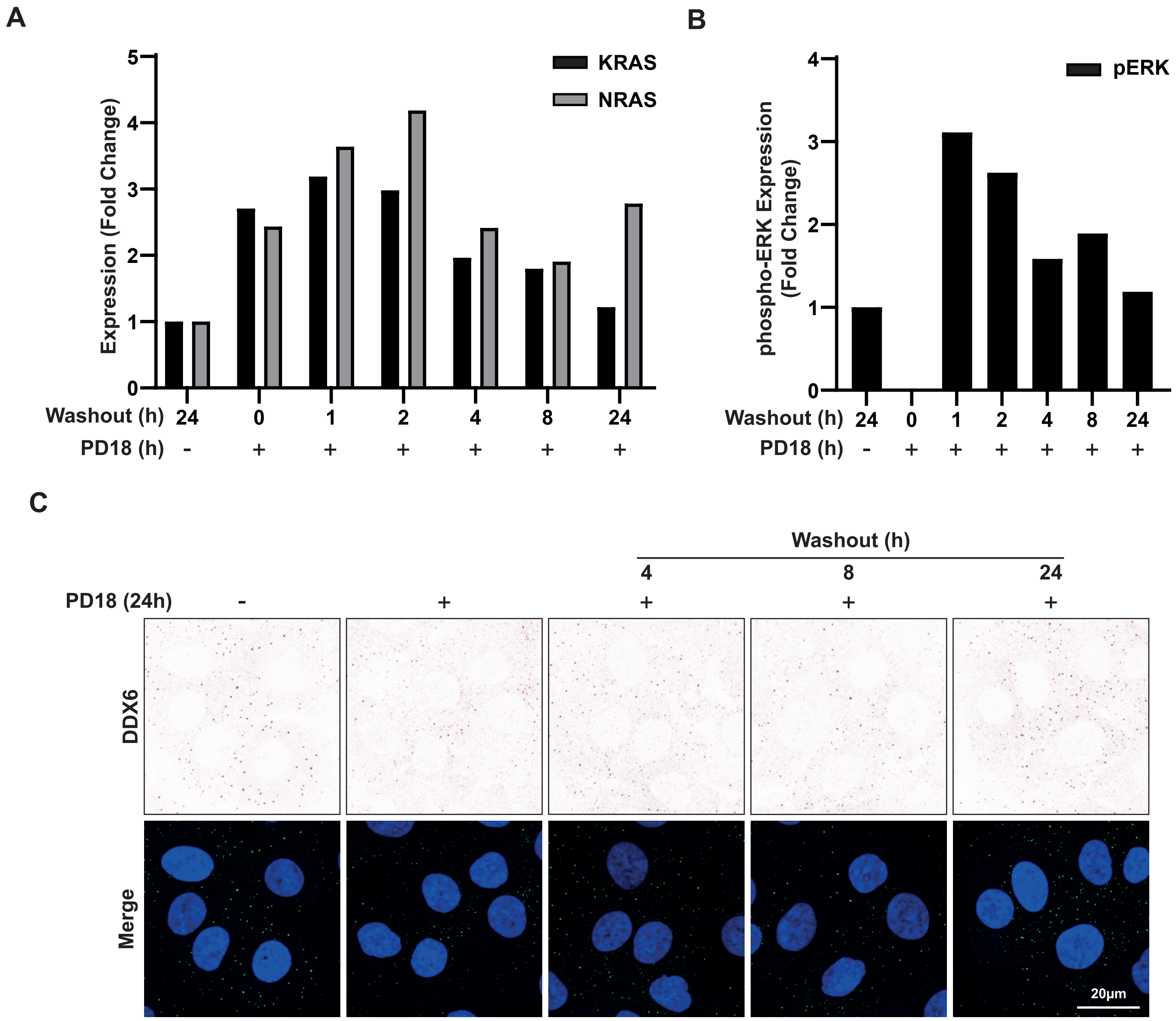


#### Supp. Figure 4

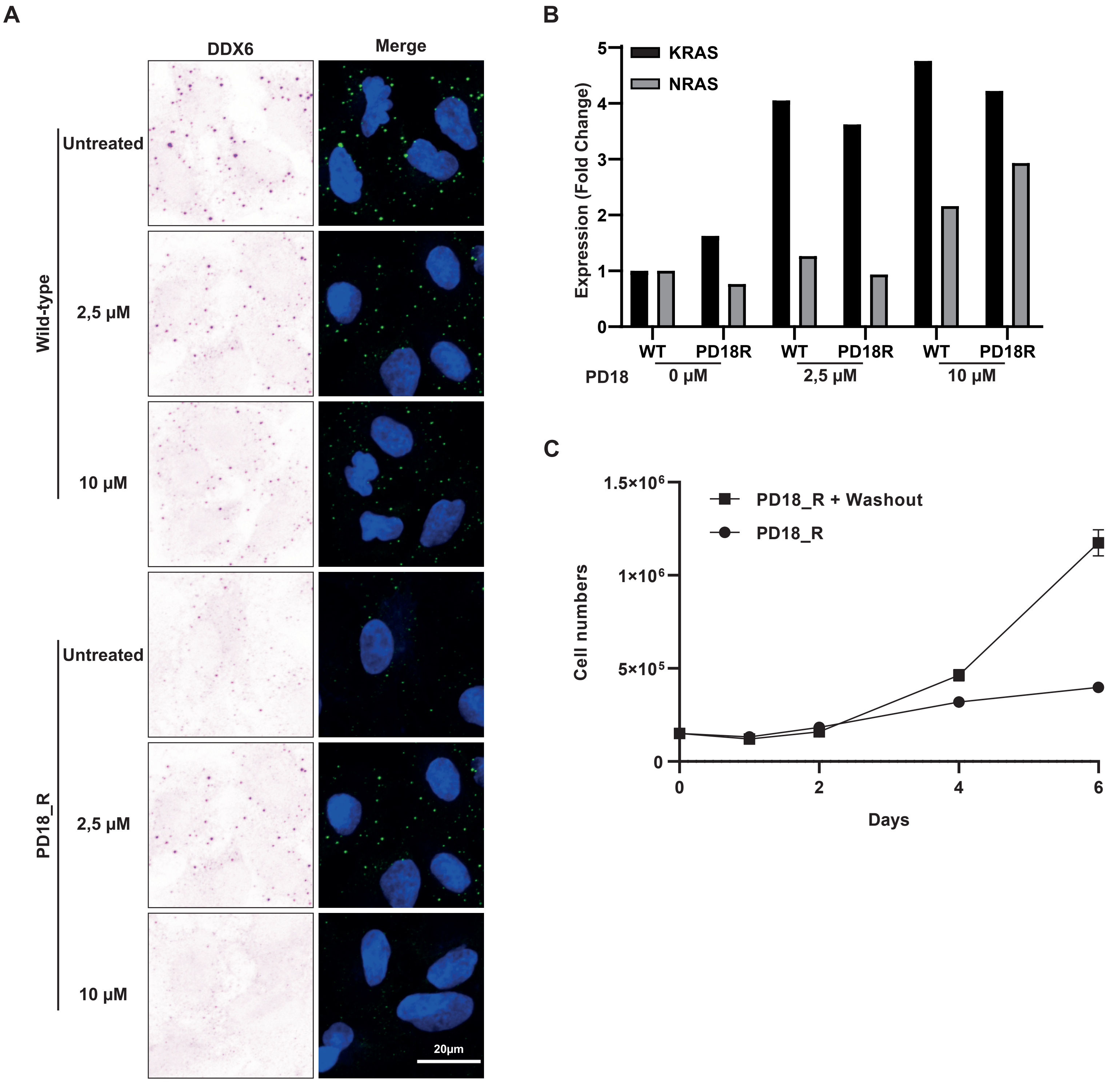


#### Supp. Figure 5

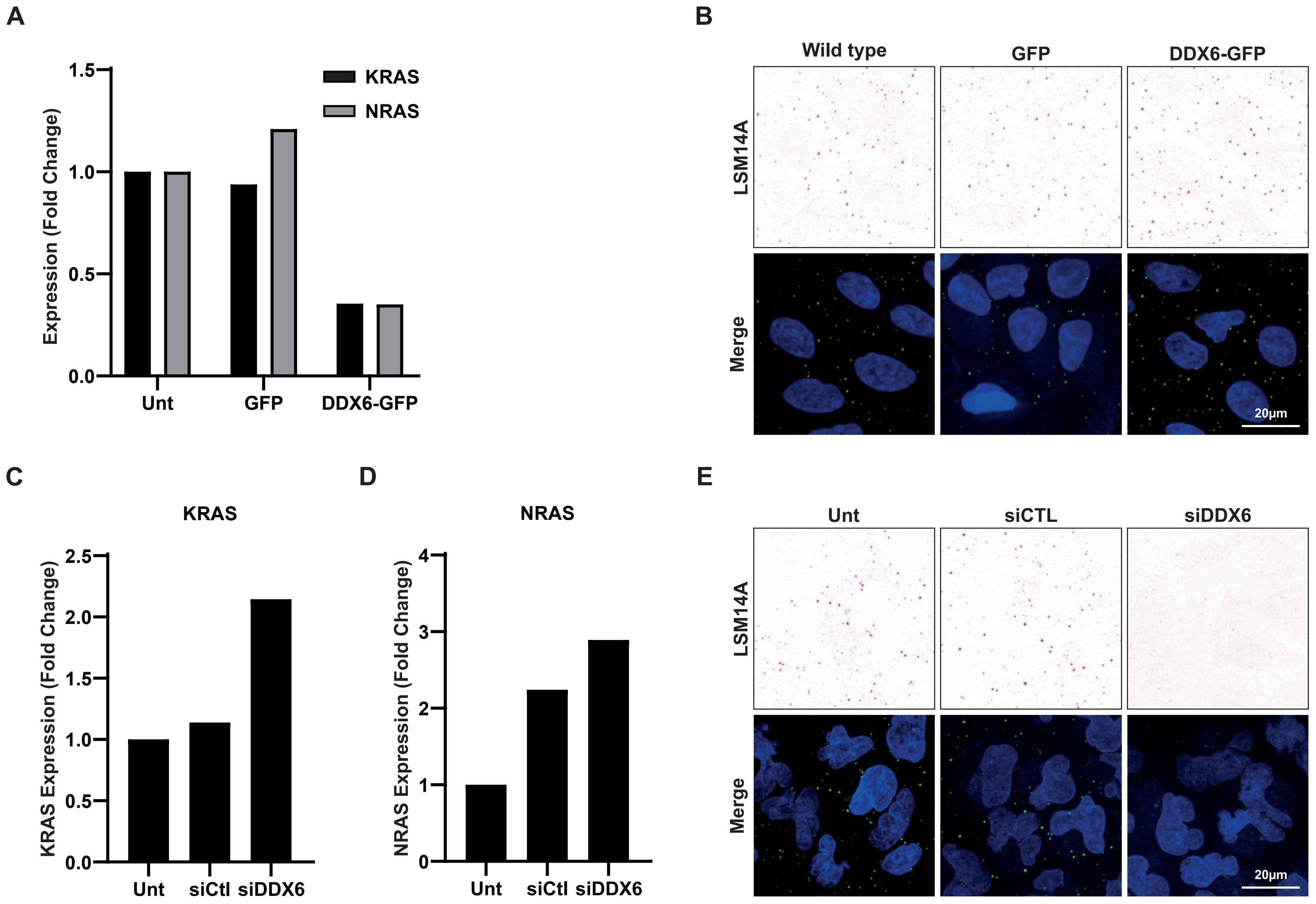


Supp. Figure 6

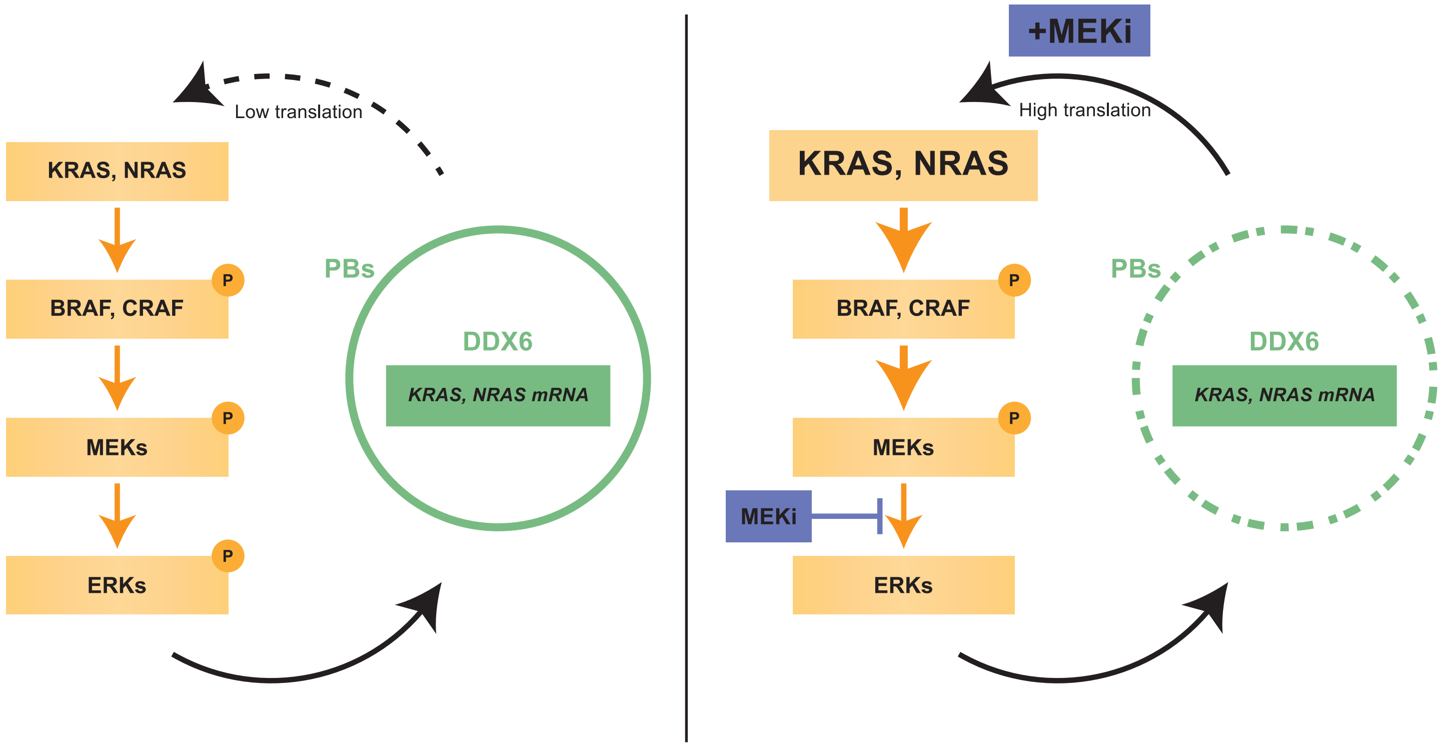
